## Supplementary information for "Engineering and implementation of synthetic molecular tools in the basidiomycete fungus *Ustilago maydis*"

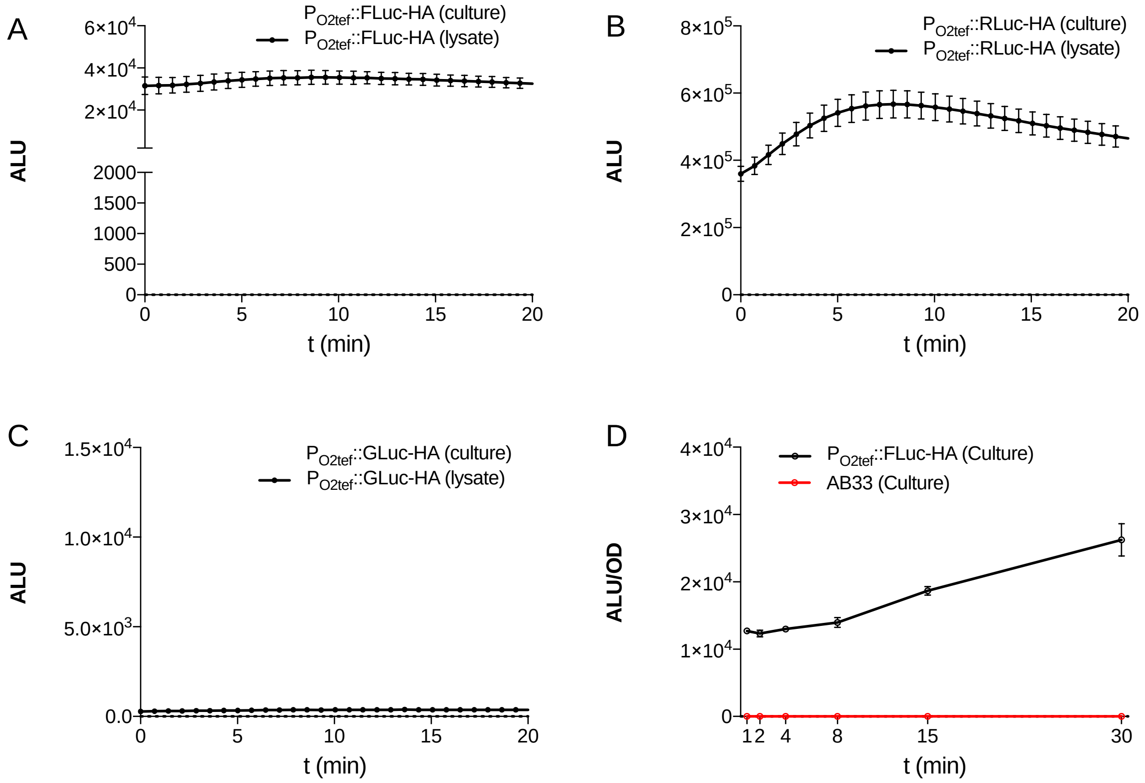


**Figure S1. Time course of luciferase activities in lysates and whole cell cultures.** Lysates of 2 ml cultures and the respective whole cell cultures, $OD_{600}$ = 0.5, of constitutively expressing luciferase strains were analyzed for their luminescence over 20 minutes after addition of substrates. FLuc (A), RLuc (B), and GLuc (C) luminescence is given in absolute luminescence units. (D) For the reporter measurement, 80 µL of the whole cell culture were transferred to 96-well assay plates and the firefly substrate was added directly and measured in 1, 2, 4, 8, 15- and 30-min. Values are normalized to an $OD_{600}$ of 0.5. Error bars represent the SEM for this individual experiment with n=3.

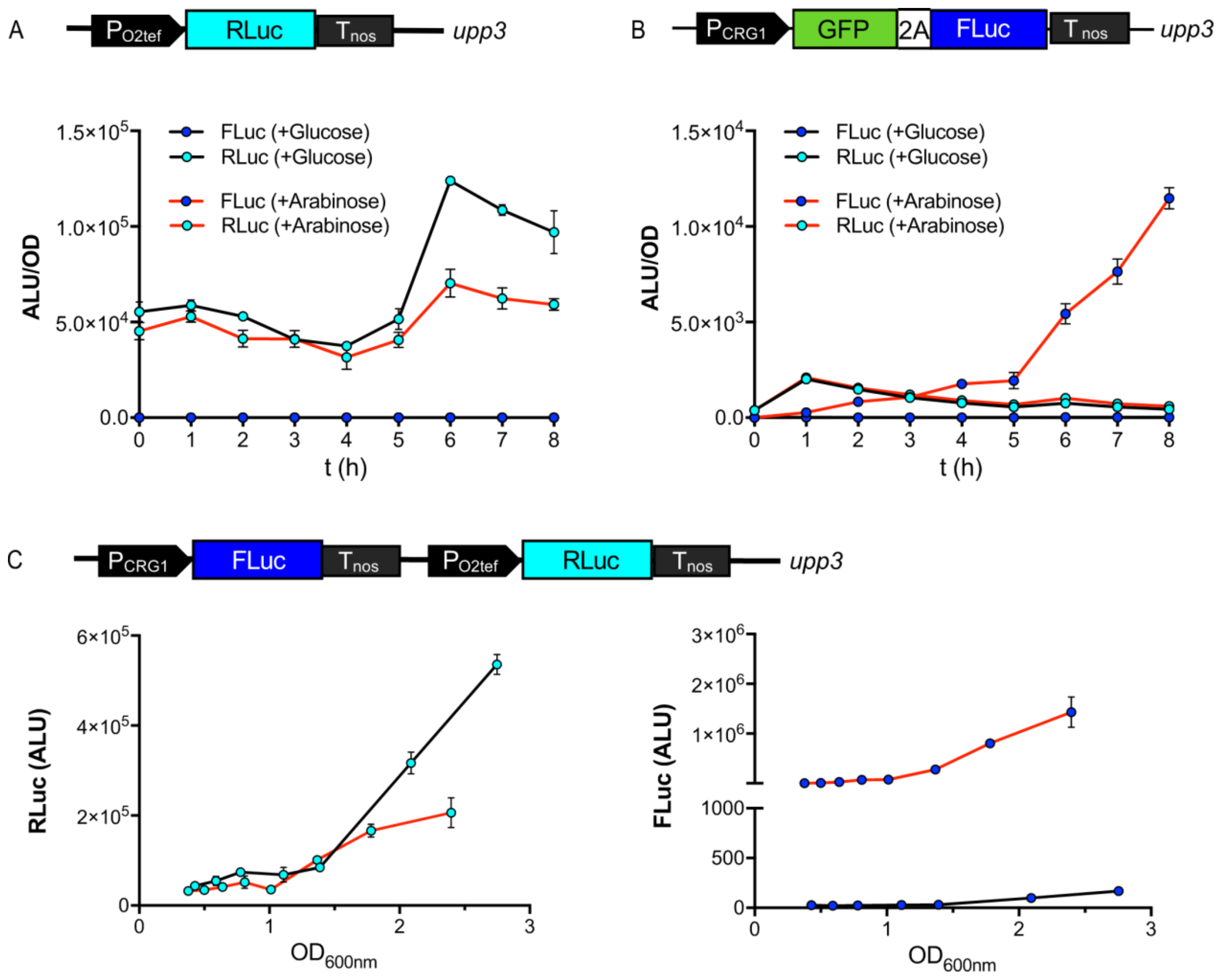


**Figure S2. Normalization of gene expression in *U. maydis*.** (A,B) single luminescence reporter strains. RLuc under control of the constitutive promotor $P_{O2tef}$ (A, sLHNH005) and FLuc (as part of a bicistronic construct) under control of the arabinose-inducible promotor $P_{crg1}$ (B, UMa3212) were cultured in CM-glucose or arabinose medium for absolute luminescence measurements. Cultures were grown overnight in CM-Glucose, at timepoint 0 determination started (black line), or cultures were shifted to CM-Arabinose (red line). 2 mL cultures were taken and lysed every hour, and analyzed for FLuc (blue dots) and RLuc (cyan dots) luminescence (normalized to an $OD_{600}$ = 0.5). (C) Dual-luminescence reporter strain (sNH039, detail information in Fig.2 and Table S1) were cultured in CM-glucose (black line) or arabinose (red line) medium. RLuc values were correlated with the $OD_{600}$ during exponential growth in both conditions, while FLuc activity only increased upon the arabinose induction.

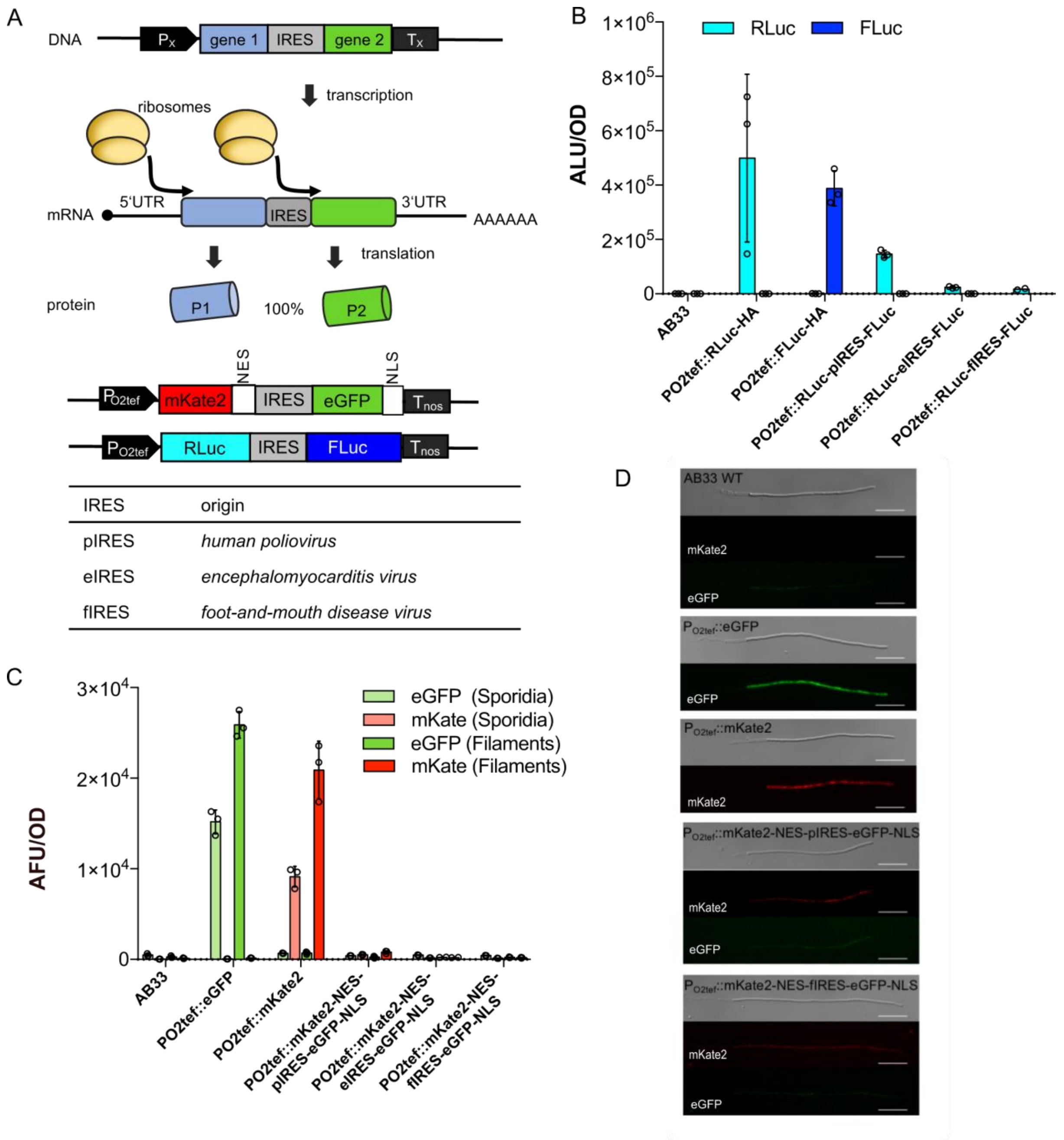


**Figure S3. Establishment of IRES sequences in *U. maydis*.** (A) Schematic representation of the IRES based bi-reporters: Genes 1 and 2 were expressed under the control of the constitutive promotor Px and are separated by IRES sequence. iRES bi-reporters strains transformed into the *upp3* locus of AB33. (B) Luminescence of whole cell lysates from 2 ml cultures, $OD_{600}$ = 0.5, of the indicated strains is shown (absolute luminescence units). (C) Fluorescence intensity of control and IRES strains measured in cultures of an $OD_{600}$ = 0.5 in sporidia, and six hours after induction of filamentous growth in a plate reader in absolute fluorescence units (AFU). In B and C, data are presented as mean values +/- SEM, n=3 individual experiments. (D) Microscopic analysis of WT AB33, mKate2, and eGFP positive controls and pIRES and fIRES strains six hours after induction of filamentous growth. Scalebars represent 20 µm.

### Table S1. Strains used in this work

| Strain | Description / Genetic background | Transformed Plasmid* | Origin |
|---|---|---|---|
| **AB33** | Induction of filamentous growth by changing the nitrogen source | | Brachmann et al., 2001 (ref13 in main text) |
| **UMa486** | AB33_IP::$P_{Otef}$-eGfp-CbxR | | Baumann et al., 2014 (ref27 in main text) (Fig.3B, S3) |
| **UMa1987** | AB33_pep4D: $P_{Otef}$-mKate2-Tnos-NatR | pUMa2986 | this work (Fig.3B, S3) |
| **UMa3212** | AB33_rrm4D_$P_{CRG}$-rrm4-gfp-linker-P2A-FLuc-HA-nosT-cbxR | | Müntjes et al., 2020 (ref15 in main text) (Fig.S2B) |
| **sLHNH005** | AB33_upp3D::$P_{O2tef}$::RLuc-HA-nosT-NatR | pLHNH030 | this work (Fig.1, 3C, S1, S2A, S3) |
| **sLHNH006** | AB33_upp3D::$P_{O2tef}$::GLuc-HA-nosT-NatR | pLHNH031 | this work (Fig.1, S1) |
| **sLHNH007** | AB33_upp3D::$P_{O2tef}$::SEAP-HA-nosT-NatR | pLHNH032 | this work (Fig.1) |
| **sLHNH008** | AB33_upp3D::$P_{O2tef}$::FLuc-HA-nosT-NatR | pLHNH033 | this work (Fig.1,3C, 4, S1, S2, S3) |
| **sNH039** | AB33_upp3D::$P_{CRG}$::FLuc-nosT-$P_{O2tef}$::RLuc-nosT-NatR | pNH054 | this work (Fig.2, S2C) |
| **sNH005** | AB33_upp3D::nosT-NES-mKate2::$P_{hCMVmin}$-CMVenhancer(5'->3')-$P_{hCMVmin}$::eGFP-NLS-nosT-NatR | pNH012 | this work (Fig. 3B) |
| **sNH006** | AB33_upp3D::nosT-NES-mKate2::$P_{hCMVmin}$-CMVenhancer(5'<-3')-$P_{hCMVmin}$::eGFP-NLS-nosT-NatR | pNH013 | this work (Fig. 3B) |
| **sNH007** | AB33_upp3D::nosT-NES-mKate2::$P_{mfa1min}$-(prf1)4(5'->3')-$P_{mfa1min}$::GFP-NLS-nosT- NatR | pNH014 | this work (Fig. 3B) |
| **sNH008** | AB33_upp3D::nosT-NES-mKate2:: $P_{mfa1min}$-(prf1)4(5'<-3')-$P_{mfa1min}$::GFP-NLS-nosT-NatR | pNH015 | this work (Fig. 3B) |
| **sNH011** | AB33_upp3D::nosT-RLuc:: $P_{hCMVmin}$-CMVenhancer(5'->3')-$P_{hCMVmin}$::FLuc-nosT-NatR | pNH030 | this work (Fig. 3C) |
| **sNH012** | AB33_upp3D::nosT-RLuc:: $P_{hCMVmin}$-CMVenhancer(5'<-3')-$P_{hCMVmin}$::FLuc-nosT-NatR | pNH031 | this work (Fig. 3C) |
| **sNH013** | AB33_upp3D::nosT-RLuc:: $P_{mfa1min}$-$(prf1)_4$(5'->3')-$P_{mfa1min}$::FLuc-nosT-NatR | pNH032 | this work (Fig. 3C) |
| **SG200** | Solopathogenic strain | | Kämper et al., 2006 (ref7 in main text) (Fig.4) |
| **SG200-FLuc (UMa3062)** | SG200_upp3D::$P_{O2tef}$::FLuc-HA-nosT-NatR | pLHNH033 | this work (Fig.4) |
| **SG200-pit1Δ** | SG200_ pit1D-HygR | | Doehlemann et al., 2011 (ref23 in main text) (Fig.4) |
| **SG200-pit1Δ-FLuc (UMa3132)** | SG200_ pit1D-HygR_upp3D::$P_{O2tef}$::FLuc-HA-nosT-NatR | pLHNH033 | this work (Fig.4) |
| **sNH001** | AB33_upp3D::$P_{O2tef}$::RLuc-pIRES-FLuc-nosT-NatR | pNH026 | this work (Fig.S3B) |
| **sNH003** | AB33_upp3D::$P_{O2tef}$::RLuc-eIRES-FLuc-nosT-NatR | pNH028 | this work (Fig.S3B) |
| **sNH004** | AB33_upp3D::$P_{O2tef}$::RLuc-fIRES-FLuc-nosT-NatR | pNH029 | this work (Fig.S3B) |
| **sLHNH009** | AB33_upp3D::$P_{O2tef}$::mKate2-NES-pIRES-eGFP-NLS-nosT-NatR | pNH009 | this work |

| | | | |
|---|---|---|---|
| | | | (Fig.S3C,D) |
| **sLHNH010** | AB33_upp3D::$P_{O2tef}$::mKate2-NES-eIRES-eGFP-NLS-nosT-NatR | pNH010 | this work (Fig.S3C,D) |
| **sLHNH011** | AB33_upp3D::$P_{O2tef}$::mKate2-NES-fIRES-eGFP-NLS-nosT-NatR | pNH011 | this work (Fig.S3C,D) |

*detail information of plasmids in Table S3

**Table S2. Generation and description of plasmids used in this work.**
All plasmids are constructed with AQUA or Gibson assembly cloning (Gibson et al., 2009; Beyer et al., 2015) if not indicated otherwise. Grey and bold: plasmids used in strain generation; white: intermediate cloning plasmids.

| Plasmid | Description | Reference |
|---|---|---|
| **pLHNH031** | **$P_{O2tef}$-GLuc-HA-nosT**<br>Vector encoding GLuc-HA under the control of $P_{O2tef}$. GLuc was amplified from pLHNH018 with oNH016 and oNH117 adding an HA-tag c-terminally to GLuc. pLHNH001 was digested with MfeI and BglII, and assembled via AUQA cloning. | this work<br>(for generating strain sLHNH006) |
| ↳ pLHNH018 | Vector encoding the synthesized codon optimized GLuc | this work |
| ↳ pLHNH001 | $P_{O2tef}$-GLuc-NLS-nosT<br>Vector encoding GLuc-NLS under the control of $P_{O2tef}$. pUMa3132 and pLHNH029 were digested with SfbI and AflII, and ligated with QuickLigase. | this work |
| ↳ ↳ pLHNH029 | Vector encoding the synthesized $P_{O2tef}$ and codon optimized GLuc-NLS | this work |
| ↳ ↳ pUMa3132 | $P_{O2tef}$-eGFP-nosT-NatR<br>Vector encoding eGFP under the control of $P_{O2tef}$ and the Nourseothricin resistance cassette for integration into the *upp3*-locus | Lee et al., 2020 (ref10 in main text) |
| **pLHNH030** | **$P_{O2tef}$-RLuc-HA-nosT**<br>Vector encoding RLuc-HA under the control of $P_{O2tef}$. pLHNH001 was digested with MfeI and BglII, RLuc was amplified from pLHNH019 with oNH020 and oNH116 adding an HA-tag c-terminally to RLuc. | this work<br>(for generating strain sLHNH005) |
| ↳ pLHNH019 | Vector encoding the synthesized codon optimized RLuc | this work |
| **pLHNH032** | **$P_{O2tef}$-SEAP-HA-nosT**<br>Vector encoding SEAP-HA under the control of $P_{O2tef}$. pLHNH001 was digested with MfeI and BglII, SEAP was amplified from pLHNH034 with oNH048 and oNH132, adding an HA-tag c-Terminally to SEAP. | this work<br>(for generating strain sLHNH007) |
| ↳ pLHNH034 | $P_{O2tef}$-SEAP-nosT<br>Vector encoding SEAP under the control of $P_{O2tef}$. pLHNH001 was digested with MfeI and BglII, SEAP n-term was amplified from pLHNH020 with oLH017 and oLH021, SEAP c-term was amplified from pLHNH035 with oNH130 and oNH131. SEAP parts were fused via PCR with oNH048 and oNH131. | this work |
| ↳ ↳ pLHNH020 | Vector encoding the synthesized codon optimized SEAP-nTerm | this work |
| ↳ ↳ pLHNH035 | Vector encoding the synthesized codon optimized SEAP-cTerm | this work |
| **pLHNH033** | **$P_{O2tef}$-FLuc-HA-nosT**<br>Vector encoding FLuc-HA under the control of $P_{O2tef}$. pLHNH001 was digested with MfeI and BglII, FLuc was amplified from pLHNH017 with oNH012 and oNH119 adding an HA-Tag c-terminally to FLuc. | this work<br>(for generating strain sLHNH008) |
| ↳ pLHNH017 | Vector encoding the synthesized codon optimized FLuc | this work |
| **pNH054** | **$P_{CGR}$-FLuc-nosT-$P_{O2tef}$-RLuc-nosT**<br>Bicistronic vector encoding FLuc under the control of the inducible $P_{CRG}$ and RLuc under the control of $P_{O2tef}$. pLHNH030 was digested with SbfI. FLuc was amplified from pLHNH017 using oLH014 and oLH018, nosT was amplified from pUMa3132 using oNH715 and oNH144. FLuc and nosT were fused via PCR using oNH717 and oNH144. $P_{CRG}$ was amplified from pUMa4175 using oNH718 and oNH719. FLuc-nosT and $P_{CRG}$ were fused via PCR using oNH720 and oNH716. | this work<br>(for generating strain sNH039) |
| ↳ pUMa4175 | $P_{CRG}$-5'UTR-rrm4-eGFP-e'UTR-nosT<br>Plasmid encoding a fusion of rrm4 and eGFP under the control of the inducible CRG promoter. | this work |

| | | |
|---|---|---|
| **pNH009** | **$P_{O2tef}$-mKate2-NES-pIRES-eGFP-NLS-nosT**<br>Bicistronic vector encoding mKate2-NES and eGFP-NLS under the control of $P_{O2tef}$. pLHNH001 was digested with MfeI and PacI, mKate2 was amplified from pLHNH015 with oNH058 and oNH122, eGFP was amplified from pLHNH015 with oNH205 and oNH057, human polio virus IRES was amplified from pKM006 with oNH124 and oNH125, mKate2, pIRES and eGFP were fused via PCR using oNH008 and oNH123. | this work<br>(for generating strain sLHNH009) |
| ↳ pKM006 | tetO13-422 bp-PhCMVmin-SEAP-pvIRES-pA<br>Vector encoding SEAP-pvIRES under the control of a modified Tet between inducible promoter | Müller et al., 2013 (ref19 in main text) |
| **pNH010** | **$P_{O2tef}$-mKate2-NES-eIRES-eGFP-NLS-nosT**<br>Bicistronic vector encoding mKate2-NES and eGFP-NLS under the control of $P_{O2tef}$. pLHNH001 was digested with MfeI and PacI, mKate2 was amplified from pLHNH015 with oNH058 and oNH122, eGFP was amplified from pLHNH015 with oNH205 and oNH057, Encephalomyocarditis virus IRES was amplified from pLHNH036 with oNH126 and oNH127, mKate2, eIRES and eGFP were fused via PCR using oNH008 and oNH123. | this work<br>(for generating strain sLHNH010) |
| ↳ pLHNH036 | Encephalomyocarditis virus IRES | this work |
| **pNH011** | **$P_{O2tef}$-mKate2-NES-fIRES-eGFP-NLS-nosT**<br>Bicistronic vector encoding mKate2-NES and eGFP-NLS under the control of $P_{O2tef}$. pLHNH001 was digested with MfeI and PacI, mKate2 was amplified from pLHNH015 with oNH058 and oNH122, eGFP was amplified from pLHNH015 with oNH205 and oNH057, Foot-and-mouthdisease virus IRES was amplified from pLHNH037 with oNH128 and oNH129, mKate2, fIRES and eGFP were fused via PCR using oNH008 and oNH123. | this work<br>(for generating strain sLHNH011) |
| ↳ pLHNH037 | Foot-and-mouthdisease virus IRES | this work |
| **pNH026** | **$P_{O2tef}$- RLuc-pIRES-FLuc-nosT**<br>Bicistronic vector encoding RLuc and FLuc under the control of $P_{O2tef}$. pLHNH001 was digested with MfeI and AscI, RLuc was amplified from pLHNH019 using oligos oLH015 and oNH181, pIRES was amplified from pKM006 using oligos oNH612 and oNH613, RLuc and pIRES were fused via PCR using oligos oNH020 and oNH613. FLuc was amplified from pLHNH017 using oligos oNH175 and oNH174. FLuc and the RLuc-pIRES fusion were fused via PCR using oligos oNH623 and oNH624. | this work<br>(for generating strain sNH001) |
| **pNH028** | **$P_{O2tef}$-RLuc-eIRES-FLuc-nosT**<br>Bicistronic vector encoding RLuc and FLuc under the control of $P_{O2tef}$. pLHNH001 was digested with MfeI and AscI, RLuc was amplified from pLHNH019 using oligos oLH015 and oNH178, eIRES was amplified from pLHNH036 using oligos oNH614 and oNH615, RLuc and eIRES were fused via PCR using oligos oNH020 and oNH615. FLuc was amplified from pLHNH017 using oligos oNH177 and oNH174. FLuc and the RLuc-IeRES fusion were fused via PCR using oligos oNH623 and oNH624. | this work<br>(for generating strain sNH003) |
| **pNH029** | **$P_{O2tef}$-RLuc-fIRES-FLuc-nosT**<br>Bicistronic vector encoding RLuc and FLuc under the control of $P_{O2tef}$. pLHNH001 was digested with MfeI and AscI, RLuc was amplified from pLHNH019 using oligos oLH015 and oNH180, fIRES was amplified from pLHNH037 using oligos oNH616 and oNH617, RLuc and fIRES were fused via PCR using oligos oNH020 and oNH617. FLuc was amplified from pLHNH017 using oligos oNH179 and oNH174. FLuc and RLuc-fIRES fusion were fused via PCR using oligos oNH623 and oNH624. | this work<br>(for generating strain sNH004) |
| **pNH012** | **nosT-NES-mKate2-$P_{hCMVmin}$-CMVenhancer(5'->3')-$P_{hCMVmin}$-eGFP-NLS-nosT** | this work |

| | | |
|---|---|---|
| | Bicistronic vector encoding mKate2-NES and eGFP-NLS under the control of a bidirectional $P_{CMV}$. pNH012a was digested with MfeI; mKate2 was amplified from pUMa2977 using oligos oNH139 and oNH141. | (for generating strain sNH005) |
| ↳ pNH012a | pNH030 (see below) was digested with AscI, eGFP was amplified from pUMa3132 using oligos oNH136 and oNH123. Parts were assembled via AQUA cloning. | this work |
| pUMa2977 | Vector containing a synthesized dicodon-optimized red fluorescent protein mKate2 with an HA-tag. | Müntjes et al., 2020 (ref15 in main text) |
| **pNH013** | **nosT-NES-mKate2-$P_{hCMVmin}$-CMVenhancer(5'<-3')-$P_{hCMVmin}$-eGFP-NLS-nosT**<br>Bicistronic vector encoding mKate2-NES and eGFP-NLS under the control of a bidirectional $P_{CMV}$. pNH013a was digested with MfeI, eGFP was amplified from pUMa3132 using oligos oNH659 and oNH660. | this work<br>(for generating strain sNH006) |
| ↳ pNH013a | pNH012 was digested with AscI, mKate2 was amplified from pUMa2977 using oligos oNH657 and oNH658. Parts were assembled via Aqua cloning. | this work |
| **pNH014** | **nosT-NES-mKate2-$P_{mfa1min}$-(prf1)$_4$(5'->3')-$P_{mfa1min}$-GFP-NLS-nosT**<br>Bicistronic vector encoding mKate2-NES and eGFP-NLS under the control of a bidirectional P(prf1)$_4$-mfa1min. pNH014a was digested with AscI, eGFP was amplified from pUMa3132 with oNH721 and oNH123. | this work<br>(for generating strain sNH007) |
| ↳ pNH014a | pNH032 was digested with MfeI; mKate2 was amplified from pUMa2977 with oNH141 and oNH153, parts were assembled via AQUA cloning. | this work |
| **pNH015** | **nosT-NES-mKate2-$P_{mfa1min}$-(prf1)4(5'<-3')-$P_{mfa1min}$-GFP-NLS-nosT**<br>Bicistronic vector encoding mKate2-NES and eGFP-NLS under the control of a bidirectional P(prf1)4-mfa1min. pNH015a was digested with AscI, mKate2 was amplified from pUMa2977 with oNH722 and oNH658. | this work<br>(for generating strain sNH008) |
| ↳ pNH015a | pNH032 was digested with MfeI, eGFP was amplified from pUMa3132 with oNH152 and oNH660. Parts were assembled via AQUA cloning. | this work |
| **pNH030** | **nosT-RLuc-$P_{hCMVmin}$-CMVenhancer(5'->3')-$P_{hCMVmin}$-FLuc-nosT**<br>Bicistronic vector encoding RLuc and FLuc under the control of a bidirectional $P_{CMV}$. pNH030a was digested with SbfI and MfeI, RLuc was amplified from pLHNH019 using oligos oLH019 and oNH196, a $P_{hCMVmin}$ was added via PCR using oligos oNH197 and oNH140, nosT was amplified from pUMa3132 using oligos oNH158 and oNH144, $P_{hCMVmin}$-RLuc and nosT were fused via PCR using oligos oNH169 and oNH143. | this work<br>(for generating strain sNH011) |
| ↳ pNH030a | pLHNH001 was digested with MfeI and AscI; FLuc was amplified from pLHNH017 using oligos oNH198 and oLH018, $P_{CMV}$ was amplified from pHB109 using oligos oNH137 and oNH135, $P_{CMV}$ and FLuc were fused via PCR using oligos oNH174 and oNH625. Parts were assembled via AQUA cloning. | this work |
| ↳ ↳ pHB109 | $P_{CMV}$-IE-PhyB-mCherry-NES-pA<br>Vector encoding PhyB-mCherry-NES under the control of the hCMV immediate early promoter. | Beyer et al., 2015 (ref22 in main text) |
| **pNH031** | **nosT-RLuc-$P_{hCMVmin}$-CMVenhancer(5'<-3')-$P_{hCMVmin}$-FLuc-nosT**<br>Bicistronic vector encoding RLuc and FLuc under the control of a bidirectional $P_{CMV}$. pNH031a was digested with MfeI. FLuc was amplified from pLHNH017 using oligos oNH184 and oNH647. | this work<br>(for generating strain sNH012) |
| ↳ pNH031a | pNH030 was digested with AscI. RLuc was amplified from pLHNH019 using oligos oNH646 and oNH645. Parts were assembled via Aqua cloning. | this work |

| | | |
|---|---|---|
| **pNH032** | **nosT-RLuc-P$_{mfa1min}$-(prf1)$_4$(5'->3')-P$_{mfa1min}$-Fluc-nosT**<br>Bicistronic vector encoding Rluc and Fluc under the control of a bidirectional P(prf1)4-mfa1min. pNH032b was digested with MfeI and SbfI. nosT was amplified from pUMa3132 with oNH144 and oNH142, Rluc was amplified from pLHNH019 with oNH627 and oNH197. nosT and Rluc were fused via PCR using oNH144 and oNH628. The resulting fragment was again amplified using oNH143 and oNH157 to add overhangs to the backbone. | this work<br>(for generating strain sNH013) |
| ↳ **pNH032b** | pNH032a was digested with MfeI and PacI, P$_{OMA}$ was amplified from pUMa2675 with oNH626 and oNH696 and parts were assembled via AQUA cloning. 4 repeats of the prf1 enhancer were lost during cloning. | this work |
| ↳ ↳ pNH032a | pLHNH001 was digested with PacI and AscI, Fluc was amplified from pLHNH017 with oNH695 and oNH174. Parts were assembled via AQUA cloning. | this work |
| ↳ ↳ pUMa2675 | P$_{OMA}$-eGFP-nosT<br>Vector encoding eGFP under the control of a prf1 operator-mfa1min promoter (P$_{OMA}$ = (prf1)8-P$_{mfa1min}$). | this work |
| **pUMa2986** | **pep4D: Potef-mKate2-Tnos-natR**<br>Vector encoding mKate2 under the control of P$_{O2tef}$ and the Nourseothricin resistance cassette for integration into the *pep4*-locus. pUMa2977 was digested with AscI and NotI to integrate mKate2 into a backbone for expression in the *pep4* locus. | this work<br>(for generating strain UMa1987) |

### Table S3. oligonucleotides used in this work.

| Oligo | Sequence (5'--3') | Description |
|---|---|---|
| oLH014 | ATGGAGGACGCCAAGAA | Fw FLuc |
| oLH015 | ATGACCAGCAAGGTCTAC | Fw RLuc |
| oLH017 | ATGGTGCTCGGTCCTT | Fw SEAP |
| oLH018 | TTAGACGGCGATCTTGC | Rev FLuc |
| oLH019 | TTACTGCTCGTTCTTGAGC | Rev RLuc |
| oLH021 | TTAGTCGATGTCCATGTTCG | Rev SEAP |
| oNH008 | CGGGATCCCCCGGGCTGCAGGAATTCGATCCCCAATTGATGGTGTCGGAGCTCAT | Fw mKate2 |
| oNH012 | CGGGATCCCCCGGGCTGCAGGAATTCGATCCCCAATTGATGGAGGACGCCAAGAA | Fw FLuc |
| oNH016 | CGGGATCCCCCGGGCTGCAGGAATTCGATCCCCAATTGATGGGCGTCAAGGTG | Fw GLuc |
| oNH020 | CGGGATCCCCCGGGCTGCAGGAATTCGATCCCCAATTGATGACCAGCAAGGTCTAC | Fw RLuc |
| oNH048 | CGGGATCCCCCGGGCTGCAGGAATTCGATCCCCAATTGATGGTGCTCGGTCCTT | Fw SEAP |
| oNH057 | CTTGTACAGCTCGTCCATG | Rev eGFP |
| oNH058 | ATGGTGTCGGAGCTCATC | Fw mKate2 |
| oNH116 | GCCGGGCGGCCGGCGCGCCGGCCGCTAGATCTTTAGGCGTAGTCGGGCACGTCGTAAGGGTAGAGCGGA<br>CCCTGCTGCTCGTTCTTGAGCAC | Rev RLuc |
| oNH117 | GCCGGGCGGCCGGCGCGCCGGCCGCTAGATCTTTAGGCGTAGTCGGGCACGTCGTAAGGGTAGAGCGGA<br>CCCTGGTCACCACCGGCAC | Rev GLuc |
| oNH119 | GCCGGGCGGCCGGCGCGCCGGCCGCTAGATCTTTAGGCGTAGTCGGGCACGTCGTAAGGGTAGAGCGGA<br>CCCTGGACGGCGATCTTGCC | Rev FLuc |
| oNH122 | CATATGGCGGTGACCG | Rev mKate2 |
| oNH123 | ATGTTTGAACGATCGCCGGGCGGCCGGCGCGCCGGCCGCTTTAGACCTTTCTCTTCTTTTTGGAGGCGCTT<br>TCTTGTACAGCTCGTCCATG | Rev eGFP |
| oNH124 | ATCTGCCCTCGAAACTCGGTCACCGCCATATGATGACCAAGAAGTTTGGCACGCTCACCATCTAGGCCGGTT<br>CGGGTGCCTCGTTAAAACAGCTCTGGGGTTG | Fw pIRES |
| oNH125 | CCGGTGAACAGCTCCTCGCCCTTGCTCACCATACAATTCGCTTTATGATAACAATCTGTGATTG | Rev pIRES |
| oNH126 | GCCGGGCGGCCGGCGCGCCGGCCGCTAGATCTTTAGGCGTAGTCGGGCACGTCGTAAGGGTAGAGCGGA<br>CCCTGCTGCTCGTTCTTGAGCAC | Rev RLuc |
| oNH127 | CGGTGAACAGCTCCTCGCCCTTGCTCACCATGATTATCATCGTGTTTTTCAAAGGAAAAC | Rev eIRES |
| oNH128 | ATCTGCCCTCGAAACTCGGTCACCGCCATATGATGACCAAGAAGTTTGGCACGCTCACCATCTAGGCCGGTT<br>CGGGTGCCTCGAGCAGGTTTCCCCAATG | Fw fIRES |
| oNH129 | CCGGTGAACAGCTCCTCGCCCTTGCTCACCATGGAAAGGAAAGGTGCCGAC | Rev fIRES |
| oNH130 | CCACCCAGCTCATCTCGAACATGGACATCGACGTCATCCTCGGTGGTG | Fw SEAP |
| oNH131 | CGCCGGGCGGCCGGCGCGCCGGCCGCTAGATCTCTATCCAGGGTGGGCG | Rev SEAP |
| oNH132 | GCCGGGCGGCCGGCGCGCCGGCCGCTAGATCTTTAGGCGTAGTCGGGCACGTCGTAAGGGTAGAGCGGA<br>CCCTGCTATCCAGGGTGGGCG | Rev SEAP |
| oNH135 | AGGCTGGATCGGTCCCGGTGTCTTCTATGGAGGTCAAAACAGCGTGGATGGCGTCTCCAGGCGATCTGAC<br>GGTTCACTAAACG | Rev PhCMVmin |
| oNH136 | CTCCATAGAAGACACCGGGACCGATCCAGCCTGGCGCGCCATGGTGAGCAAGGGCG | Fw GFP |
| oNH137 | TATTAATAGTAATCAATTACGGGGTCATTAGTTC | Fw GFP |
| oNH139 | CACGCTGTTTTGACCTCCATAGAAGACACCGGGACCGATCCAGCCTCAATTGATGGTGTCGGAGCTCATC | Fw mKate2 |
| oNH140 | AATGGGCGGTAGGCGTGTACGGTGGGAGGTCTATATAAGCAGAGCTCGTTTAGTGAACCGTCAGATCGCC<br>TGGAGACGCCATCCACGCTGTTTTGACCTCC | Fw PhCMV |
| oNH141 | CCAAATGTTTGAACGATCGCCGGGCGGCCCAATTGCTAGATGGTGAGCGTGCCAAACTTCTTGGTCATCAT<br>ATGGCGGTGACCG | Rev mKate2 |
| oNH142 | GGCCGCCCGG | |
| oNH143 | TCACCATAGCAGGCCTAGATGGCCCCTGCAGGCTCATGTTTGACAGCTTATCATCG | Fw nosT |
| oNH144 | CTCATGTTTGACAGCTTATCATCG | Rev nosT |
| oNH152 | TTGAACATCAAATCAACTACCTTACTCTATCACAATTGATGGTGAGCAAGGGCG | Fw eGFP |
| oNH153 | TTGAACATCAAATCAACTACCTTACTCTATCACAATTGATGGTGTCGGAGCTCATC | Fw mKate2 |
| oNH157 | AGTGTGGCACTCGAATCCCCCTGCTCGAGAAGAATCCGACAGCCAAACCTC | Fw Pmfa1min |
| oNH158 | GCCCGGCGATCGTTC | Fw nosT |
| oNH169 | ACTAGTCAATAATCAATGTCAACATGGCGGTCCAAATGGGCGGTAGGCG | Fw PhCMVmin |
| oNH174 | TTGCCAAATGTTTGAACGATCGCCGGGCGGCCGGCGCGCCGGCCGCTTTAGACGGCGATCTTGC | Rev FLuc |
| oNH175 | ACAATCACAGATTGTTATCATAAAGCGAATTGGCGATCGCATGGAGGACGCCAAGAA | Fw FLuc |
| oNH177 | ACGTGGTTTTCCTTTGAAAAACACGATGATAAGCGATCGCATGGAGGACGCCAAGAA | Fw FLuc |
| oNH178 | TCAAGAAGACAGGGCCAGGTTTCCGGGCCCTCGCGATCGCTTACTGCTCGTTCTTGAGC | Rev RLuc |
| oNH179 | AATAGGTGACCGGAGGTCGGCACCTTTCCTTTGCGATCGCATGGAGGACGCCAAGAA | Fw FLuc |
| oNH180 | TTGCACGTTTTGTGTCATTGGGGAAACCTGCTGCGATCGCTTACTGCTCGTTCTTGAGC | Rev RLuc |
| oNH181 | CTGGGGTGGGTACAACCCCAGAGCTGTTTTAAGCGATCGCTTACTGCTCGTTCTTGAGC | Rev RLuc |
| oNH184 | CTCCATAGAAGACACCGGGACCGATCCAGCCTCAATTGATGGAGGACGCCAAGAA | Fw FLuc |
| oNH196 | CACGCTGTTTTGACCTCCATAGAAGACACCGGGACCGATCCAGCCTCAATTGATGACCAGCAAGGTCTAC | Fw RLuc |
| oNH197 | TTGCCAAATGTTTGAACGATCGCCGGGCGGCCCAATTGTTACTGCTCGTTCTTGAGC | Rev RLuc |
| oNH198 | CTCCATAGAAGACACCGGGACCGATCCAGCCTGGCGCGCCATGGAGGACGCCAAGAA | Fw FLuc |
| oNH205 | ATGGTGAGCAAGGGCG | Fw mCerulean,mVenus, mCherry,eGFP |

| | | |
|---|---|---|
| oNH612 | TTAAAACAGCTCTGGGGTTG | Fw pIRES |
| oNH613 | CAATTCGCTTTATGATAACAATCTGTGATTG | Rev pIRES |
| oNH614 | GAGGGCCCGGAAAC | Fw eIRES |
| oNH615 | TTATCATCGTGTTTTTCAAAGGAAAAC | Rev eIRES |
| oNH616 | AGCAGGTTTCCCCAATG | Fw fIRES |
| oNH617 | AAAGGAAAGGTGCCGAC | Rev fIRES |
| oNH623 | CGGGATCCCCCGG | Fw IRES-fusion |
| oNH624 | TTGCCAAATGTTTGAACGATCG | Rev IRES-fusion |
| oNH625 | CGGGATCCCCCGGGCTGCAGGAATTCGATCCCCAATTGGACCGCCATGTTGACATTGATTATTGACTAGTTA TTAATAGTAATCAATTACGGGGTCATTAGTTC | Fw $P_{CMV}$ |
| oNH626 | CGGGATCCCCCGGGCTGCAGGAATTCGATCCCCAATTGCTTCTCGAGCAGGGGG | Fw $P_{OMA}$ |
| oNH627 | TCACTTCTCGCCCGTTCTTTTGAACATCAAATCAACTACCTTACTCTATCACAATTGATGACCAGCAAGGTCT AC | Fw RLuc |
| oNH628 | AATCCGACAGCCAAACCTCATCCACTCTCACTTTCACACTCTAACTTATACGATCACTTCTCGCCCGTTC | Fw Pmfa1min |
| oNH645 | TTGCCAAATGTTTGAACGATCGCCGGGCGGCCGGCGCGCCGGCCGCTTTACTGCTCGTTCTTGAGC | Rev RLuc |
| oNH646 | CTCCATAGAAGACACCGGGACCGATCCAGCCTGGCGCGCCATGACCAGCAAGGTCTAC | Fw RLuc |
| oNH647 | TTGCCAAATGTTTGAACGATCGCCGGGCGGCCCAATTGTTAGACGGCGATCTTGC | Rev FLuc |
| oNH657 | CTCCATAGAAGACACCGGGACCGATCCAGCCTGGCGCGCCATGGTGTCGGAGCTCATC | Fw mKate2 |
| oNH658 | TGTTTGAACGATCGCCGGGCGGCCGGCGCGCCCTAGATGGTGAGCGTGCCAAACTTCTTGGTCATCATATG GCGGTGACCG | Rev mKate2 |
| oNH659 | CTCCATAGAAGACACCGGGACCGATCCAGCCTCAATTGATGGTGAGCAAGGGC | Fw eGFP |
| oNH660 | TTGCCAAATGTTTGAACGATCGCCGGGCGGCCCAATTGGGCCGCTTTAGACCTTTCTCTTCTTTTTTGGAGG CGCTTTCTTGTACAGCTCGTCCATG | Rev eGFP |
| oNH695 | AAGATCAAGGGTGCCGGTGGTGACTAATTAATTAAAACTTCTCGCCCGTTCTTTTGAACATCAAATCAACTA CCTTACTCTATCAGGCGCGCCATGGAGGACGCCAAGAA | Fw FLuc |
| oNH696 | TGATTTGATGTTCAAAAGAACGGGCGAGAAGTGATCGTATAAGTTAGAGTGTGAAAGTGAGAGTGGATGA GGTTTGGCTGTCGGATTCTCCCTTATATCCTTGACGGTAC | Rev POMA |
| oNH715 | AGGCCAAGAAGGGTGGCAAGATCGCCGTCTAAGCGATCGCGGCCGCCCGG | Fw nosT |
| oNH716 | CGAGCTCGGTACGGGGGATCCACTAGTTCTAGCTCATGTTTGACAGCTTATCATCG | Rev nosT |
| oNH717 | GAGGCCAAAAAAGATACCATAATAGGCCTGAGTTAATTAAATGGAGGACGCCAAGAA | Fw FLuc |
| oNH718 | CATAGTACATCAGGCTACTAACTGTC | Fw $P_{CRG}$ |
| oNH719 | CTCAGGCCTATTATGGTATCTTTTTTG | Rev $P_{CRG}$ |
| oNH720 | CACGCGTCTCACCATAGCAGGCCTAGATGGCCCCTGCAGGCATAGTA | Fw $P_{CRG}$ |
| oNH721 | TTGAACATCAAATCAACTACCTTACTCTATCAGGCGCGCCATGGTGAGCAAGGGCG | Fw eGFP |
| oNH722 | TTGAACATCAAATCAACTACCTTACTCTATCAGGCGCGCCATGGTGTCGGAGCTCATC | Fw mKate2 |
